# ATG5 regulates autophagy-apoptosis-ER stress dysregulation in steroid-induced osteonecrosis of the femoral head (SONFH) pathogenesis

**DOI:** 10.64898/2026.03.07.710256

**Authors:** Kunkun Liu, Boyong Jiang, Wanlin Liu, Yulin Gong, Zhenqun Zhao

**Affiliations:** The Second Affiliated Hospital of Inner Mongolia Medical University, Hohhot, 010030, China; Traumatic Surgery, Beijing Jishuitan Hospital,Capital Medical University, Beijing,100035, China; The Affiliated Hospital of Inner Mongolia Medical University, Hohhot, 010020, China

**Author notes:** Corresponding author. E-mail addresses, Mailing address: No. 59, Horqin South Road, Saihan District, Hohhot, Inner Mongolia, China. These authors contributed equally to this work (Kunkun Liu, Boyong Jiang).

**Keywords:** Steroid-induced osteonecrosis of the femoral head, ATG5, Autophagy, Endoplasmic reticulum stress, Apoptosis

## Abstract

The aim is to investigate how ATG5 regulates autophagy, endoplasmic reticulum stress (ERS), and apoptosis in steroid-induced osteonecrosis of the femoral head (SONFH), and to evaluate ATG5-targeted inhibition as a SONFH intervention. Differentially expressed genes in SONFH were screened using GEO dataset GSE74089. Autophagy/apoptosis/ERS pathway activities were analyzed via GSEA, GO/KEGG enrichment, and GSVA. Rat bone marrow mesenchymal stem cells (BMSCs) were isolated for osteoblast modeling. Groups included: control (A), steroid-treated (B, methylprednisolone), and intervention (C, steroid + ATG5-siRNA). Autophagosome formation, apoptosis rate, and ATG5/PERK/LC3 expression were assessed by electron microscopy, flow cytometry, qPCR, and western blotting. Sixty SD rats were divided into three groups (as above). The SONFH model was established via intramuscular methylprednisolone injection, with intervention group receiving ATG5-siRNA. Bone pathology, cell death, and pathway regulation were evaluated using HE/TUNEL staining, electron microscopy, and molecular detection. Transcriptome analysis revealed synergistic activation of autophagy (ATG5/BECN1), apoptosis (CASP3/9/12), and ERS (PERK) pathways in SONFH, with ATG5 strongly correlating with all three. Steroids upregulated ATG5 to overactivate autophagy, triggering PERK-mediated ERS and ERS-specific apoptosis. ATG5-siRNA intervention inhibited autophagosome formation, reduced apoptosis, downregulated PERK, and alleviated trabecular fractures and empty lacunae. ATG5 deficiency blocked PERK signaling, suppressing both autophagic death and ERS-dependent apoptosis.ATG5 drives autophagy overactivation and ERS-apoptosis cascades via PERK pathway activation, constituting a core SONFH mechanism. Targeted ATG5 silencing effectively blocks this pathology, offering novel preventive/therapeutic strategies.

## 1. Introduction

Osteonecrosis of the femoral head (ONFH) is a disabling disorder characterized by femoral head ischemia and cell death, causing structural and functional bone deterioration [1–3]. ONFH etiology includes traumatic and non-traumatic forms [4], with steroid use, chronic alcohol intake, and trauma history as established risk factors [5]. Steroid-induced ONFH (SONFH) predominates in non-traumatic cases [6], primarily affecting patients receiving excessive glucocorticoids (GCs) for immune-related conditions [7,8].

GCs contribute crucially to SONFH pathogenesis. They induce lipid metabolism disorders causing intraosseous hypertension, impair femoral head perfusion, and promote necrosis [9]. Additionally, GCs directly trigger apoptosis in bone-forming osteoblasts and maintenance osteocytes [10], accelerating bone loss. GCs also dysregulate autophagy [11], potentially compromising osteocyte function and increasing ONFH susceptibility. GC exposure promotes ferroptosis, exacerbating bone loss and SONFH progression [12]. Crucially, GCs activate endoplasmic reticulum stress (ERS), damaging osteogenesis and angiogenesis—key disease mechanisms [13].

Autophagy is a lysosomal degradation process for intracellular components (proteins, lipids, organelles) [14]. Triggered primarily by nutrient deprivation, it also occurs during differentiation, development, or organelle damage [15]. Autophagosomes fuse with lysosomes forming autolysosomes, where contents undergo proteolytic degradation [16]. While basal autophagy maintains cellular homeostasis, its upregulation primarily occurs during physiological remodeling or pathological stress [17]. Excessive autophagy degrades normal organelles alongside damaged ones, causing autophagic cell death (Type II programmed death) [18]. This non-apoptotic death mode significantly contributes to developmental differentiation and remodeling [19].

Although GC-induced autophagy dysregulation, apoptosis, and ERS in SONFH are partially characterized, key regulators and their crosstalk mechanisms require elucidation. This study aims to delineate ATG5’s role in SONFH pathogenesis using integrated transcriptomic, cellular, and animal approaches. These findings may provide new insights into molecular mechanisms converting aberrant autophagy to autophagic death in SONFH, while establishing an experimental foundation for ATG5-targeted therapies.

## 2. Method

### Analysis workflow of GEO database

We retrieved the GEO dataset (GSE74089) using the GEOquery package in R. Using the limma package, we screened differentially expressed genes (DEGs) to identify significant differences between SONFH and normal samples. Expression heterogeneity across samples was subsequently analyzed. We performed functional enrichment analyses (including GSEA) using clusterProfiler, focusing on pathway activation status related to apoptosis execution, autophagy regulation, and endoplasmic reticulum stress response. GO enrichment explored biological processes involving DEGs, while KEGG pathway analysis identified enriched signaling pathways. Using the GSVA package, we evaluated activity differences in autophagy regulation, endoplasmic reticulum stress, and apoptosis-related functions between SONFH and control groups. The corrplot package analyzed expression correlations among autophagy, apoptosis execution, and key endoplasmic reticulum stress genes, emphasizing association strengths between core genes. Finally, we validated core DEG expression levels using dataset samples.

### Isolation and culture of BMSCs

Three-week-old SD rats were anesthetized, sacrificed, and immersed in 75% ethanol (Aladdin Biochemical Technology, Shanghai, China) for 30 minutes. We harvested hind limb bones in a clean bench (Suzhou Clean Air Technology, Suzhou, China), rinsed bone marrow cavities with DMEM, then seeded cell suspensions into culture flasks and maintained them at 37°C with 5% CO₂. Initial medium change occurred at 16 hours. At 80% confluence, we digested cells with trypsin and passaged them at 1:3 ratio, using fourth-passage cells for experiments.

### Osteogenic induction and identification of BMSCs

Fourth-passage BMSCs underwent 21-day culture in osteogenic induction medium (containing hormones, ascorbic acid, and β-glycerophosphate sodium). After 95% ethanol fixation (Aladdin Biochemical Technology, Shanghai, China), we performed alizarin red staining for 30 minutes and observed osteogenesis microscopically (Olympus, Tokyo, Japan).

### Construction and pretreatment of osteoblast model

We seeded osteogenically induced cells (5×10⁴ cells/well) in 24-well plates divided into: Group A (control), Group B (SONFH), and Group C (SONFH + siRNA-ATG5)(Genomeditech, Shanghai, China). After cationic liposome transfection, we administered drugs and collected samples at 12-72 hours post-administration.

### Experimental animals and establishment of SONFH model

Animal grouping: Sixty clean-grade SD rats were randomized into three groups (n=20). Group A received intramuscular saline (2 ml/rat, three doses at 24-hour intervals). Group B received methylprednisolone (20 mg/kg; Nanjing Saihongrui Biotechnology, Nanjing, China) identically. Group C received identical methylprednisolone injections plus 0.5 mol/kg ATG5-siRNA (Genomeditech, Shanghai, China) via cationic liposomes.

### Staged sampling

Weekly from 1-4 weeks post-modeling, we sacrificed five rats per group and harvested bilateral femoral heads. Samples were: fixed in 4% paraformaldehyde (for HE/TUNEL/immunofluorescence); fixed in 2.5% glutaraldehyde (Sinopharm Chemical Reagent, Shanghai, China) for EM; or stored at -80°C (Aucma, Qingdao, China) for qRT-PCR/WB.

### Transmission electron microscopy for autophagosomes

After trypsin digestion and centrifugation, cells were fixed in 2.5% glutaraldehyde, post-fixed in osmium tetroxide (Zhenzhun Biotechnology, Shanghai, China), gradient-dehydrated, embedded in resin (Solarbio, Beijing, China), sectioned, stained, and imaged.

### CCK-8 viability assay

Cells seeded in 96-well plates received CCK-8 solution. After 4-hour incubation, we measured OD450nm to calculate viability.

### Cellular immunofluorescence

Cells fixed with 4% paraformaldehyde were permeabilized with 0.1% Triton X-100, blocked with 5% BSA (Solarbio, Beijing, China), incubated with primary antibody → fluorescent secondary antibody (Abcam, Cambridge, UK) → DAPI counterstain, then imaged (Leica, Shanghai, China).

### Flow cytometry for apoptosis

After trypsin digestion and PBS washing (Shengxing Biotechnology, Nanjing, China), cells were stained with Annexin V-APC/PI and analyzed by flow cytometry.

### HE staining

Following dewaxing and hydration, sections underwent hematoxylin staining (Solarbio, Beijing, China) for 10 minutes, hydrochloric acid-alcohol differentiation (Nanjing Chemical Reagent, Nanjing, China), bluing, and eosin counterstaining (Beyotime, Shanghai, China). After mounting, we observed trabecular bone and empty lacunae by light microscopy (Olympus, Tokyo, Japan).

### TUNEL staining

After dewaxing, Proteinase K permeabilization and blocking, we applied DNase I/TdT enzyme reaction solution (Beyotime, Shanghai, China), developed color with DAB, and counterstained with hematoxylin (Solarbio, Beijing, China) to detect apoptosis.

### Electron microscopy detection

For autophagosome observation, 0.2 cm subchondral tissues were decalcified, fixed in osmium tetroxide (Zhenzhun Biotechnology, Shanghai, China), sectioned, and stained. We observed ultrastructure by scanning electron microscopy (Hitachi, Tokyo, Japan) after dehydration, drying, and mounting.

### Tissue immunofluorescence

After dewaxing and antigen retrieval, we blocked endogenous peroxidase with 3% H₂O₂ (Sinopharm Chemical Reagent, Shanghai, China) and non-specific binding with 5% BSA (Solarbio, Beijing, China). Sections were incubated with primary antibody (4°C overnight) → fluorescent secondary antibody (Abcam, Cambridge, UK) → DAPI counterstain, then imaged for target expression.

### RT-PCR

Cells cultured in CO₂ incubators (Thermo Fisher, Waltham, USA) reached 70-90% confluence before PBS washing (Shengxing Biotechnology, Nanjing, China). After 0.25% trypsin digestion (HyClone, Logan, USA) and timely termination, we pelleted cells by centrifugation and seeded them in 6-well plates (Costar, Washington, USA) at 1×10⁵ cells/well. Total RNA extraction used TRIzol (Takara, Shiga, Japan): chloroform phase separation, isopropanol precipitation, ethanol washing, and dissolution in DEPC water (Aladdin Biochemical Technology, Shanghai, China). We measured A260/A280 ratios by spectrophotometry (Thermo Fisher, Waltham, USA), removed genomic DNA, and synthesized cDNA using PrimeScript RT kits (Takara, Shiga, Japan). qPCR was performed in quantitative PCR systems (Applied Biosystems, California, USA) with SYBR® Premix Ex Taq (Takara, Shiga, Japan).

### Western Blot

Exosomes lysed in 2% SDS buffer (ice, 30 min) were centrifuged at 12,000×g. Protein concentrations were determined by BCA assay (Beyotime, Shanghai, China). After SDS-PAGE electrophoresis (Bio-Rad, California, USA) using 30% polyacrylamide gels (Solarbio, Beijing, China), proteins were electrotransferred to PVDF membranes (Millipore, Burlington, USA) using transfer systems (Bio-Rad, California, USA). For immunodetection: membranes blocked with 5% skim milk (BD, New Jersey, USA); incubated overnight (4°C) with CD63/CD81 primary antibodies (Abcam, Cambridge, UK); incubated (2 h, RT) with secondary antibodies (Abcam, Cambridge, UK); developed using chemiluminescence systems (Tanon, Shanghai, China) with ECL substrate (Tanon, Shanghai, China).

### Statistical analysis

Data are expressed as mean ± SD and analyzed using GraphPad Prism 6. We employed t-tests for two-group comparisons and ANOVA for multi-group comparisons, considering P < 0.05 statistically significant.

## 3. Result

### Differential gene analysis revealed that dysregulation of the autophagy-apoptosis-endoplasmic reticulum (ER) stress network is linked to the pathological mechanism of steroid-induced osteonecrosis of the femoral head (SONFH)

Analysis of GEO transcriptome data revealed significant gene expression differences between SONFH and normal samples. Differentially expressed genes (DEGs) included NFH, CASP12, BECN1, CASP9, CASP3, ATG5, and EIF2AK3 (PERK) (Fig. 1a, c), with marked inter-sample heterogeneity in expression levels (Fig. 1d).

**Fig 1.**
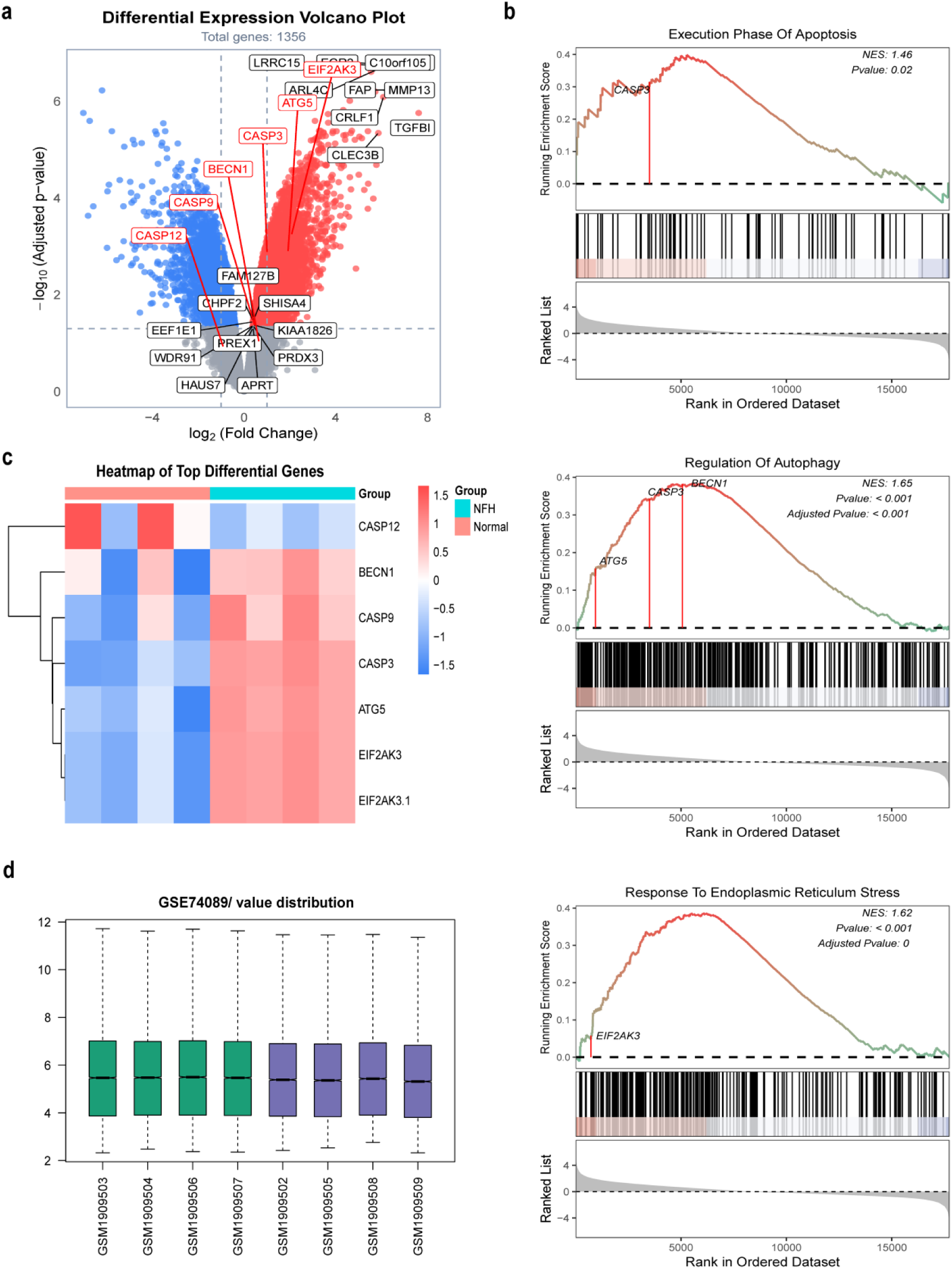
a. Volcano plot of differentially expressed genes b. Results of gene set enrichment analysis (GSEA) c. Heatmap of top differentially expressed genes d. Box plot

Gene Set Enrichment Analysis (GSEA) showed that pathways related to the apoptotic execution phase, autophagy regulation (involving BECN1, CASP3, ATG5, etc.), and ER stress response (involving EIF2AK3) were significantly activated or regulated in SONFH (Fig. 1b). GO and KEGG enrichment analyses confirmed that DEGs were significantly enriched in biological processes (e.g., bone development, collagen fiber organization, MHC class II protein complex assembly) and pathways (e.g., focal adhesion, PI3K-Akt, autophagy, TGF-β). Among these, bone development-related genes (e.g., COL1A1, OGN) were predominantly downregulated, while MHC class II assembly genes (e.g., HLA family) were predominantly upregulated, indicating that SONFH involves bone matrix metabolic disorders, abnormal collagen structure, and immune response activation (Fig. 2a-d).

**Fig 2.**
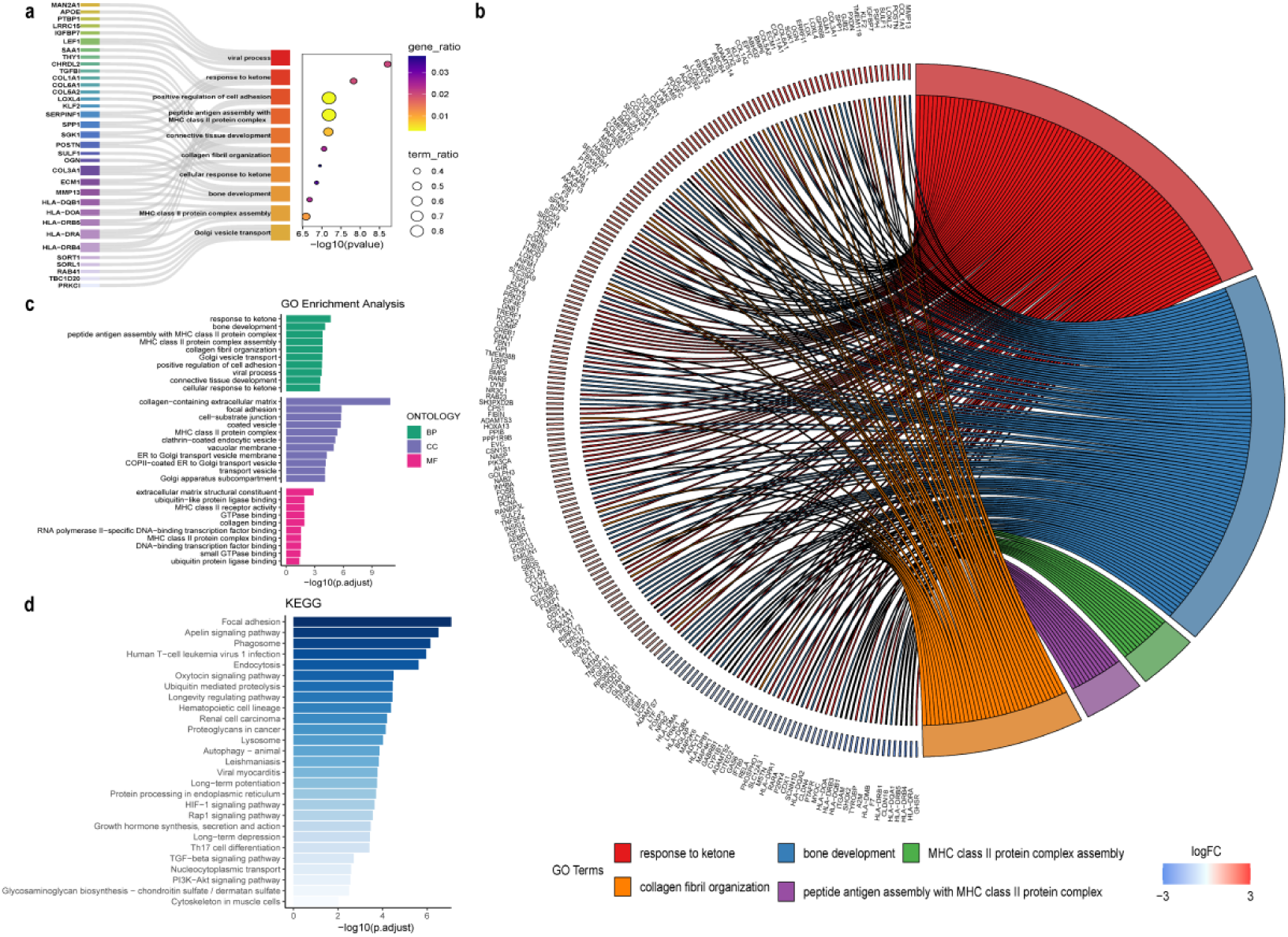
a-b. Results of genes corresponding to GO enrichment analysis c. Results of the top 10 GO enrichment analyses d. Results of KEGG pathway enrichment analysis

GSVA analysis indicated that autophagy regulation, ER stress, and apoptotic functions were significantly enhanced in the SONFH group (Fig. 3a). Autophagy genes (ATG5, ATG12, BECN1), apoptotic execution genes (CASP3/9/12), and the key ER stress gene EIF2AK3 (PERK) showed extensive correlations; notably, ATG5 exhibited strong positive correlations with all three (Fig. 3b, d). Differential expression of core genes validated these findings (Fig. 3c). In summary, abnormal networks involving activated autophagy, apoptosis, and ER stress, together with bone matrix metabolic disorders and immune activation, constitute key pathological mechanisms of SONFH.

**Fig 3.**
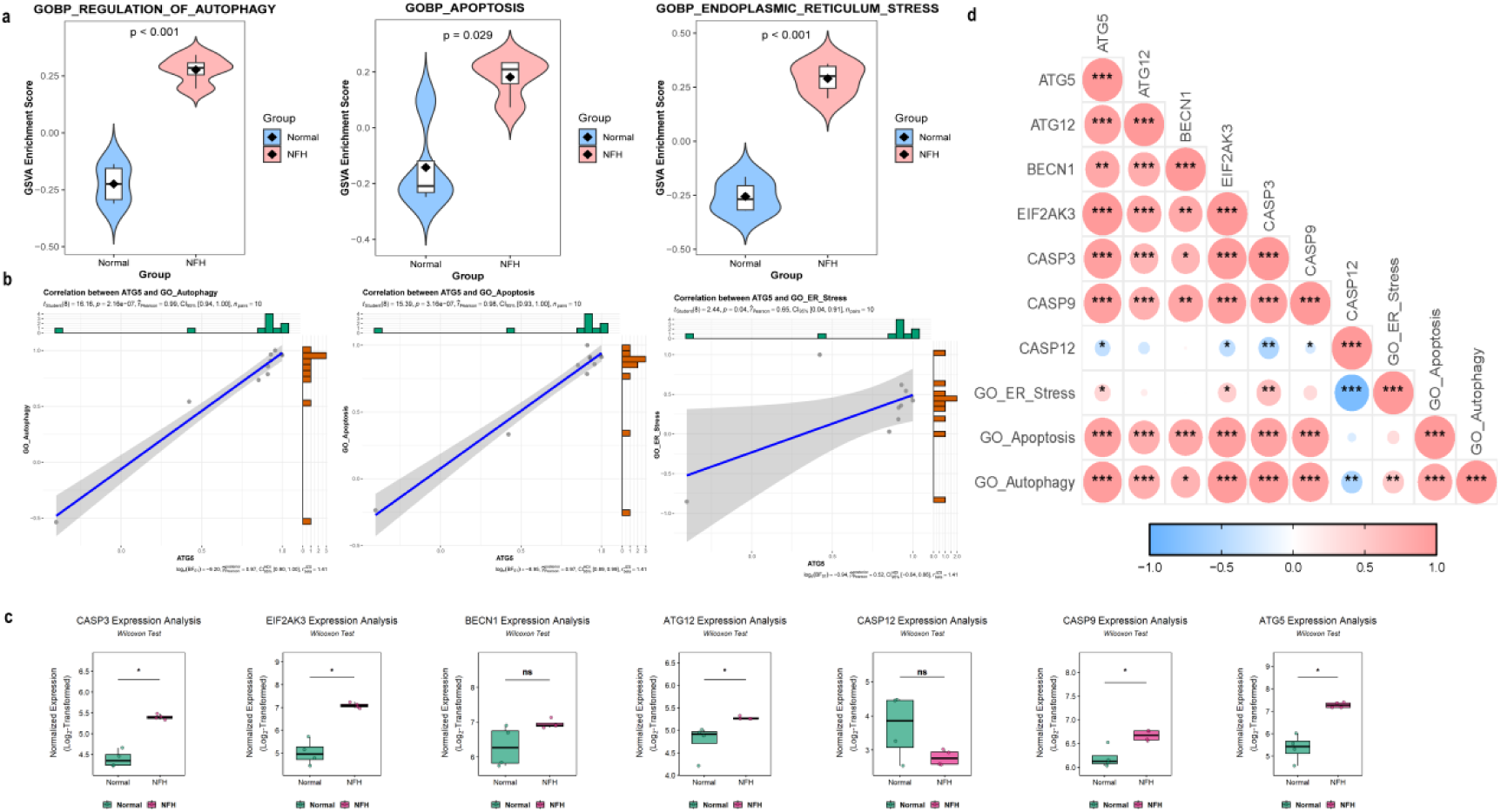
a. GSEA score results b. Correlation analysis of ATG5 with endoplasmic reticulum stress, apoptosis, and autophagy c. Expression levels of core genes d. Results of correlation analysis between molecules

### Cellular experimental results

#### Isolation, culture, and osteogenic differentiation of rat BMSCs

Rat bone marrow mesenchymal stem cells (BMSCs) were isolated using the bone marrow adherence method, and high-purity primary cells were successfully obtained via gradient centrifugation combined with differential adherence selection. Under conventional culture conditions, these cells exhibited a typical spindle or polygonal morphology, with mononuclear characteristics, grew in a monolayer adherent manner, and conformed to the typical stem cell biological properties of BMSCs. To verify their osteogenic differentiation potential, after 21 days of osteogenic induction culture, the cell morphology changed significantly: in addition to retaining some spindle-shaped characteristics, the cells more frequently showed cord-like arrangement, polygonal aggregated growth, and a distinct trend of stratified growth, indicating that the cells had entered the osteogenic differentiation stage. Further detection via Alizarin Red staining revealed the formation of numerous deep red mineralized nodules in the extracellular matrix. These nodules, as characteristic markers of calcium salt deposition secreted by osteoblasts, fully confirmed that rat BMSCs had successfully differentiated into osteoblasts (Fig. 1-a).

### Hormones activate autophagy via the ATG5-dependent pathway

To investigate the regulatory effect of hormones on autophagy, transmission electron microscopy was used in this study to observe changes in the number and structure of autophagosomes. The results showed that in Group B (hormone-treated group), the number of autophagosomes increased significantly with prolonged treatment time, and the autophagosomes had intact structures with clearly encapsulated substances to be degraded (Fig. 4-d). In contrast, in Group C (hormone + siRNA-ATG5 intervention group), the number of autophagosomes was significantly lower than that in Group B, and the structural integrity of some autophagosomes was reduced, with phenomena such as blurred membrane structures and leakage of contents. Further molecular-level detection results showed that in Group B, the mRNA transcription level and protein expression of the key autophagy gene ATG5 were continuously upregulated over time; meanwhile, the autophagy marker indices, including the LC3-II/LC3-I ratio and Beclin-1 protein expression, increased synchronously, indicating enhanced autophagic activity. In Group C, after specific intervention with siRNA-ATG5, the basal expression of ATG5 was significantly inhibited, the hormone-induced upregulation trend of ATG5 was completely blocked, and the above autophagy-related indices were all significantly suppressed. These results confirm that hormones can activate the formation and maturation of autophagosomes via the ATG5-dependent pathway. Furthermore, correlation analysis between autophagy levels and the endoplasmic reticulum stress marker PERK revealed a positive feedback regulatory relationship between them: increased PERK expression could promote the upregulation of autophagy levels, and enhanced autophagic activity, in turn, promoted PERK expression (Fig. 4-f, Fig. 5-a, b).

**Fig 4.**
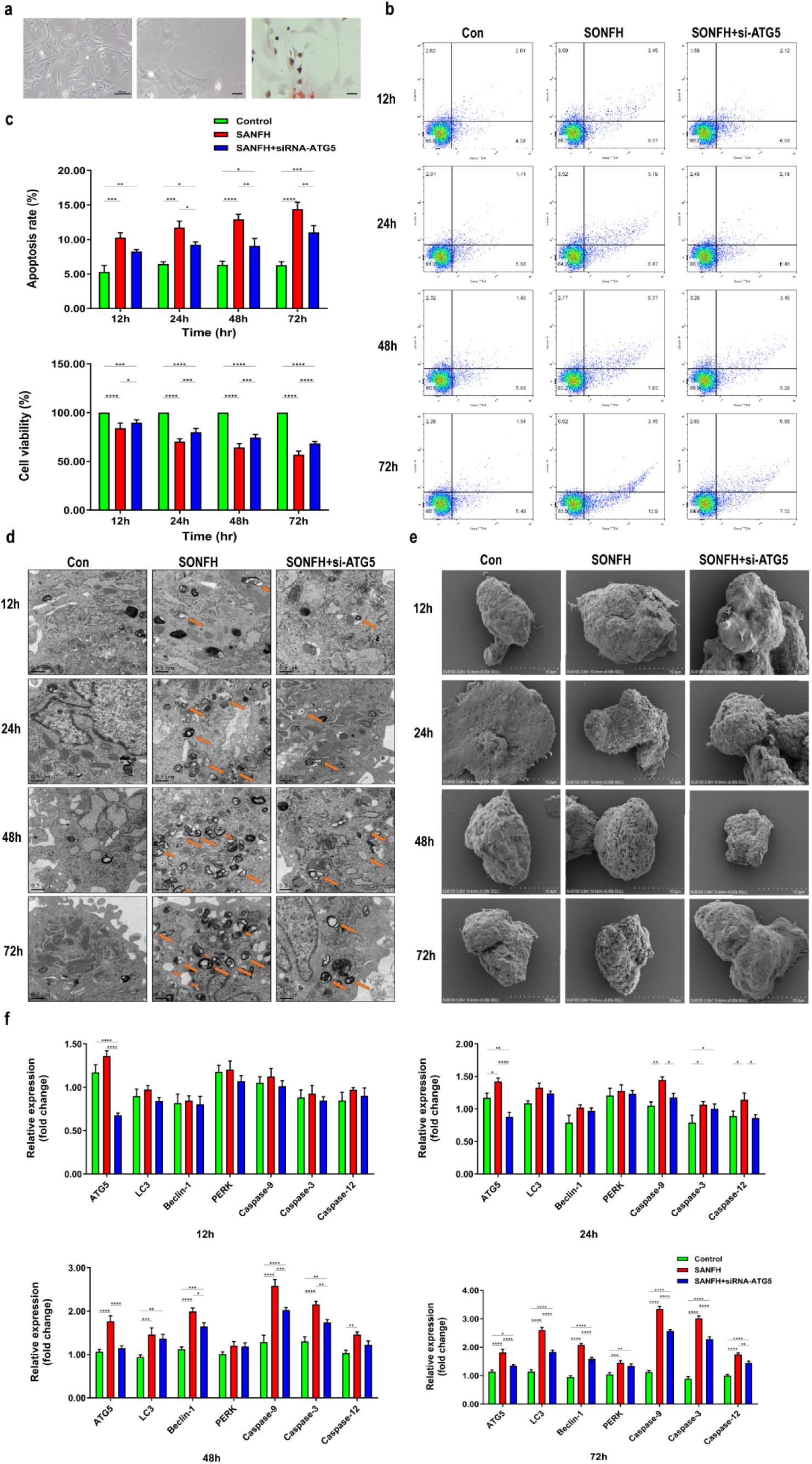
a. Morphology of bone marrow mesenchymal stem cells under an inverted microscope (100x, 200×) and the morphology of cells after BMSCs induction stained with Alizarin Red (100×) (scale bar, 100 μm) b. Flow cytometry apoptosis plots of osteoblasts in each group at 12 h, 24 h, 48 h, and 72 h c. Statistics on the proportion of cell apoptosis and comparison of the survival rates of osteoblasts in each group at 12 h, 24 h, 48 h, and 72 h d. Observation of the ultrastructure of osteoblasts under an electron microscope (20,000×) e. Observation of the ultrastructure of cells in each group under a scanning electron microscope (1,000×) f. Detection of mRNA expression levels of related molecules by fluorescence quantitative PCR at 12 h, 24 h, 48 h, and 72 h (* P < 0.05, ** P < 0.01, *** P < 0.001, **** P < 0.0001 vs. Control)

**Fig 5.**
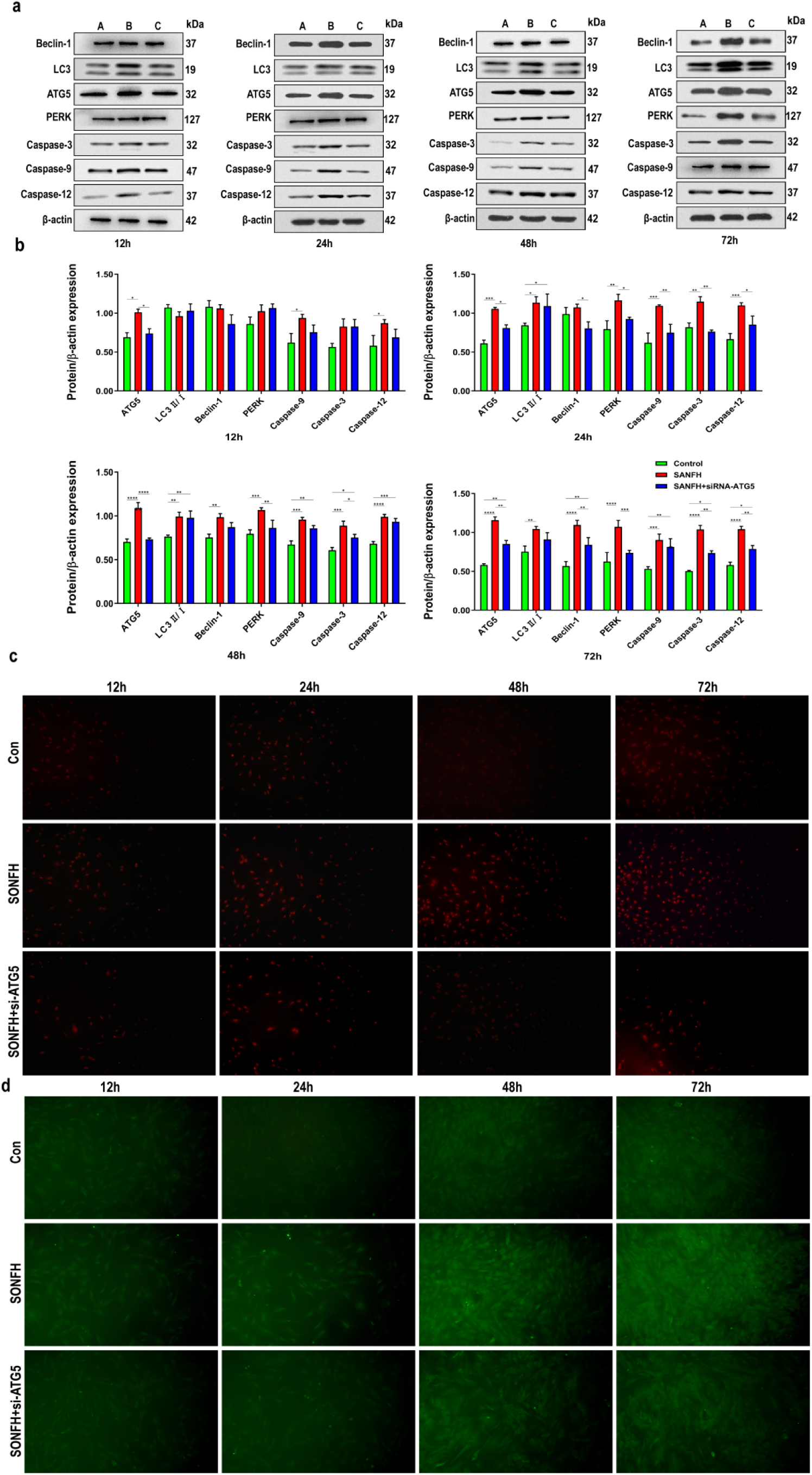
a-b. Comparison of protein expression levels of related molecules in each group detected by Western blot at 12 h, 24 h, 48 h, and 72 h (* P < 0.05, ** P < 0.01, *** P < 0.001, **** P < 0.0001 vs. Control) c. Immunofluorescence detection of ATG5 protein expression in each group at 12 h, 24 h, 48 h, and 72 h d. Immunofluorescence detection of Beclin-1 protein expression in each group at 12 h, 24 h, 48 h, and 72 h

### Effects of hormones on cell survival and apoptosis and the role of ATG5

To clarify the effects of hormones on cell survival and apoptosis and the role of ATG5 therein, the CCK-8 assay was used in this study to detect cell viability. The results showed that in Group B (SONFH model group) treated with methylprednisolone, cell viability decreased in a time-dependent manner with prolonged treatment, with the most significant decrease compared to the control group at 72 h of treatment. In contrast, cell viability in Group C (ATG5 knockdown group) was significantly higher than that in Group B at the same time points, suggesting that ATG5 deficiency could partially reverse the cytotoxicity induced by hormones. Further results from flow cytometry-based apoptosis analysis showed that the apoptosis rate in Group B gradually increased with prolonged treatment time, mainly consisting of late apoptosis. The apoptosis rate in Group C was significantly lower than that in Group B, with a also significant decrease in the proportion of late apoptosis (Fig. 4-b, c, e). These results indicate that hormones can induce cell apoptosis in a time-dependent manner, while ATG5 silencing can partially alleviate hormone-induced apoptosis, exerting a certain protective effect on cells (Fig. 4-b, c, e). Combined results from the two experiments reveal that hormones induce cell apoptosis in a time-dependent manner, and ATG5 plays a pro-apoptotic role in this process; its silencing can protect cells by reducing apoptosis.

### Crosstalk among the endoplasmic reticulum stress-autophagy-apoptosis molecular network

To reveal the crosstalk among the molecular networks of endoplasmic reticulum stress, autophagy, and apoptosis, multi-dimensional detection was performed in this study. The results showed that in Group B (hormone-treated group), PERK, a key molecule in the endoplasmic reticulum stress pathway, was continuously activated, with its phosphorylation level increasing significantly over time. Meanwhile, the expression levels of autophagy markers (LC3-II/LC3-I, Beclin-1) and apoptosis-related factors (especially Caspase-12, specific to the endoplasmic reticulum stress PERK pathway) increased synchronously, suggesting a possible synergistic activation relationship among the three. In Group C (ATG5-silenced group), ATG5 silencing not only inhibited the process of autophagic flux but also significantly downregulated the expression level of PERK and the activity of Caspase family members (including Caspase-3, Caspase-9, and Caspase-12). These results indicate that ATG5 deficiency can block the transmission of PERK signals, thereby synergistically inhibiting overactivated autophagic activity and endoplasmic reticulum stress-specific apoptosis, suggesting that ATG5 is a key node connecting the endoplasmic reticulum stress-autophagy-apoptosis molecular network (Fig. 4-f, Fig. 5 a-d, Fig. 6 a-e).

**Fig 6.**
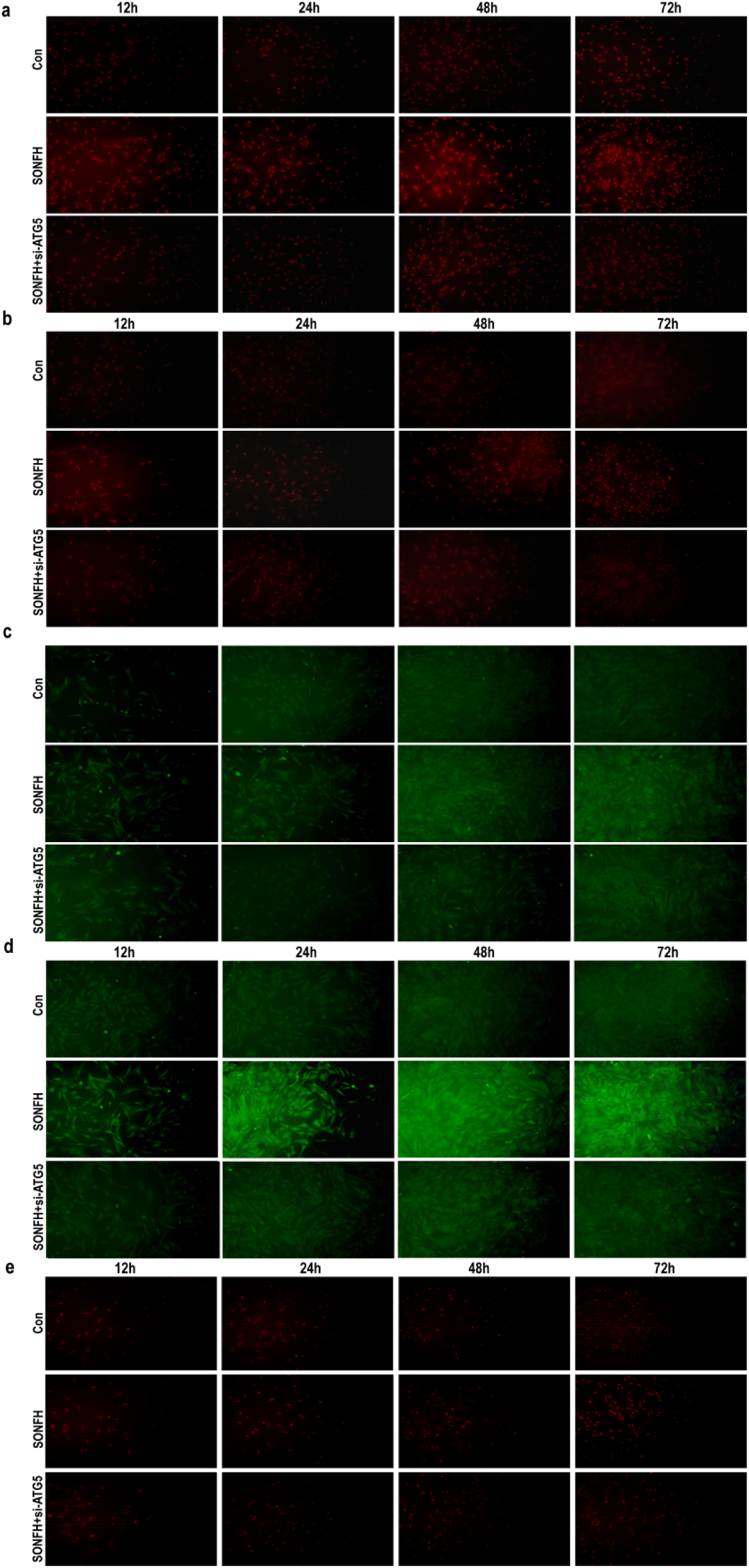
a. Immunofluorescence detection of Caspase-3 protein expression in each group at 12 h, 24 h, 48 h, and 72 h b. Immunofluorescence detection of Caspase-9 protein expression in each group at 12 h, 24 h, 48 h, and 72 h c. Immunofluorescence detection of Caspase-12 protein expression in each group at 12 h, 24 h, 48 h, and 72 h d. Immunofluorescence detection of LC3 protein expression in each group at 12 h, 24 h, 48 h, and 72 h e. Immunofluorescence detection of PERK protein expression in each group at 12 h, 24 h, 48 h, and 72 h

### Animal experimental results

#### Hormone-induced pathological damage to bone tissue and the protective effect of ATG5-siRNA

To observe hormone-induced pathological damage to bone tissue and the intervention effect of ATG5-siRNA, bone tissue morphology was analyzed in this study using HE staining and scanning electron microscopy. The results showed that in Group B (hormone-treated group), the trabecular bone was disorderly arranged with irregular orientation, accompanied by multiple fractures; the trabecular spacing was increased, the number of empty bone lacunae was significantly increased, and bone structure collapse was observed in some areas. In Group C (hormone + ATG5-siRNA group), the number of trabecular fractures was significantly reduced, the arrangement was more regular than that in Group B, the number of empty bone lacunae was significantly lower than that in Group B, and the integrity of the bone structure was significantly improved. The results of scanning electron microscopy were consistent with those of HE staining: the surface of trabecular bone in Group B was rough and uneven with numerous fracture gaps, while that in Group C was relatively smooth with fewer fracture gaps, suggesting that ATG5-siRNA exerts a significant protective effect against hormone-induced bone tissue damage (Fig. 7-a, d).

**Fig 7.**
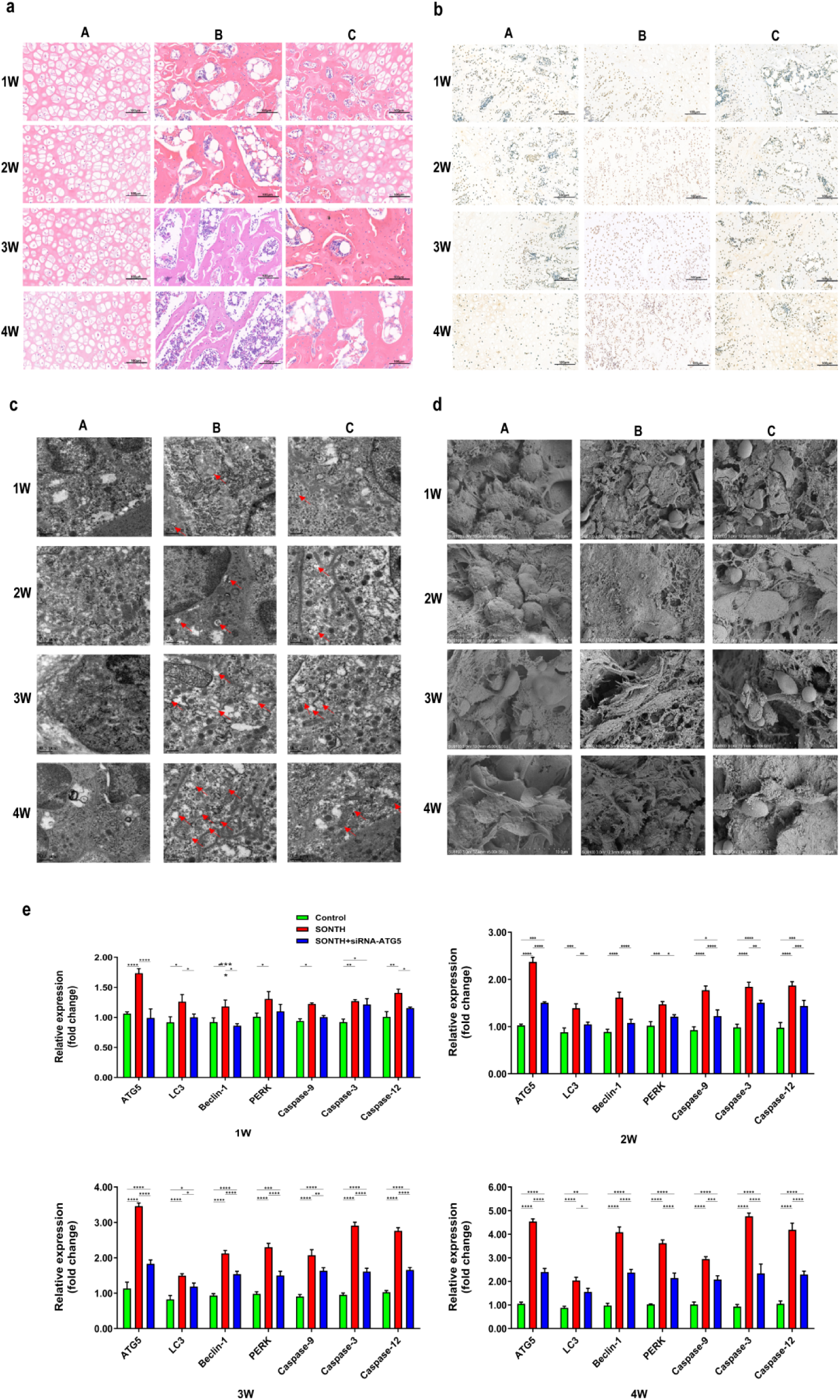
a. Results of HE staining (200×, n = 3) b. Results of TUNEL staining (100×, n = 3) c. Observation of the ultrastructure of femoral head tissue in each group of rats under a transmission electron microscope (10,000×). The red arrows in the figure indicate autophagosomes. d. Observation of the ultrastructure of femoral head tissue in each group of rats under a scanning electron microscope (1,000×) e. Comparison of mRNA expression levels of related molecules after 1 W, 2 W, 3 W, and 4 W of animal modeling (* P < 0.05, ** P < 0.01, *** P < 0.001, **** P < 0.0001 vs. Control)

### Hormone-induced autophagy activation and conversion to autophagic cell death

By observing via transmission electron microscopy and detecting autophagy markers, this study investigated the effect of hormones on autophagic activity in bone tissue and the intervention effect of ATG5-siRNA. The results showed that in Group B (hormone group), the number of autophagosomes in bone tissue cells showed an increasing trend with prolonged treatment time: it slightly increased compared to the control group at 1 week, significantly increased at 2 weeks, and peaked at 3-4 weeks. At this time, some cells exhibited excessive autophagy activation, characterized by the accumulation of numerous autophagosomes, excessive degradation of organelles, and eventual conversion to autophagic cell death. Molecular-level detection showed that the expression of autophagy markers such as ATG5, Beclin-1, and LC3 in Group B increased continuously over time, consistent with the changing trend in the number of autophagosomes. In Group C (hormone + ATG5-siRNA group), the above indices were significantly reduced: the number of autophagosomes at each time point was lower than that in Group B, and the expression levels of ATG5, Beclin-1, and LC3 were also significantly downregulated. These findings indicate that ATG5-siRNA can effectively block hormone-induced autophagic processes and inhibit the conversion of excessive autophagy activation to autophagic cell death (Fig. 7-c, e; Fig. 8 a-d).

**Fig 8.**
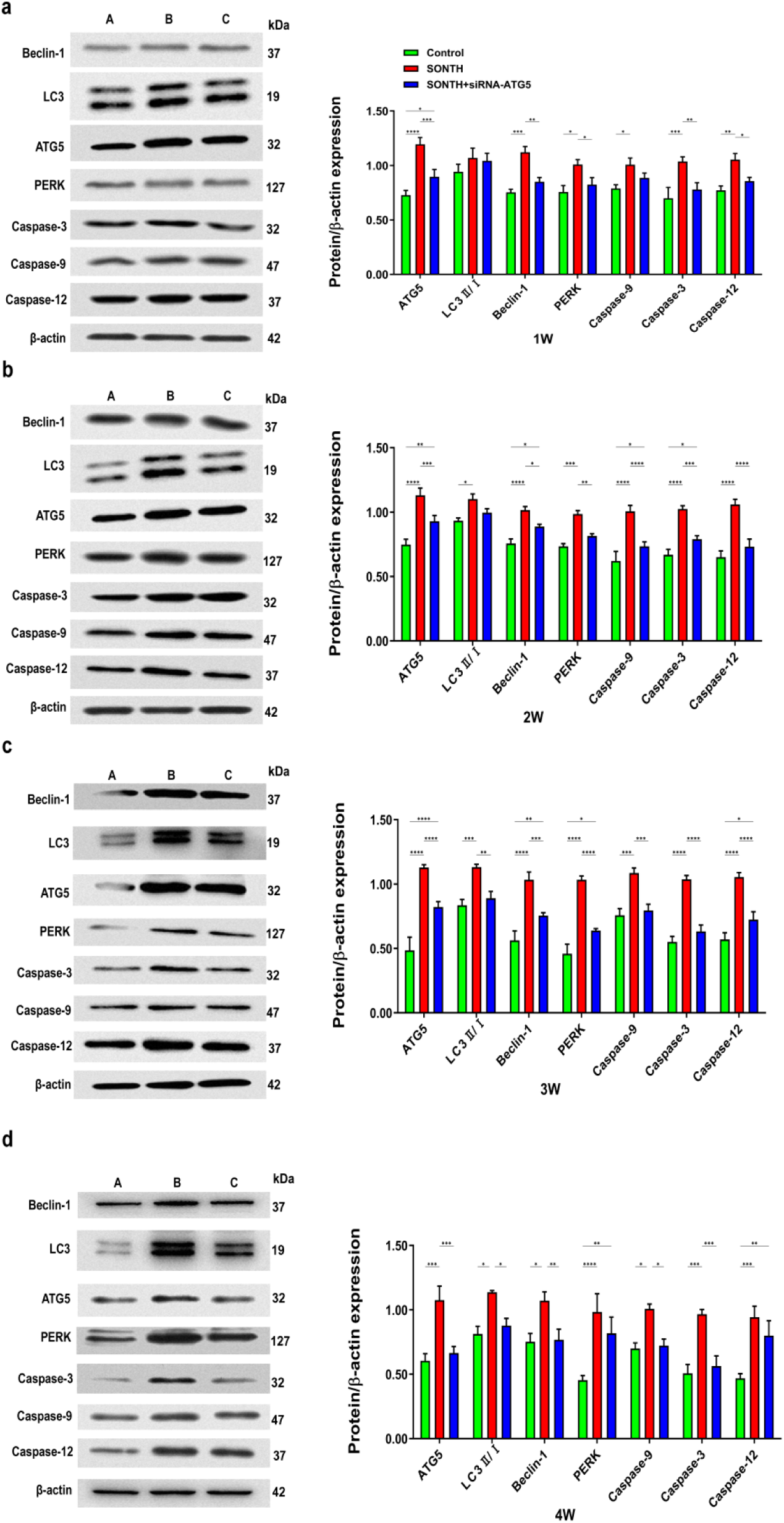
a-d. Comparison of protein expression levels of related molecules after 1 W, 2 W, and 3 W of animal modeling (* P < 0.05, ** P < 0.01, *** P < 0.001, **** P < 0.0001 vs. Control)

### Hormone activation of the endoplasmic reticulum stress-apoptosis pathway and the inhibitory effect of ATG5-siRNA

To clarify the effect of hormones on apoptosis of bone tissue cells and the underlying mechanism, analysis was performed in this study using TUNEL staining and detection of apoptosis-related factors. TUNEL staining results showed that the number of apoptotic cells in bone tissue in Group B (hormone group) showed an increasing trend with prolonged treatment time. Molecular-level detection showed that the expression of Caspase-3, Caspase-9, Caspase-12, and the endoplasmic reticulum stress marker PERK in Group B was significantly increased, with a changing trend consistent with that of cell apoptosis. In Group C (hormone + ATG5-siRNA group), the number of apoptotic cells at the same time points was significantly lower than that in Group B, and the expression levels of Caspase-3/9/12 and PERK were also significantly lower than those in Group B. These results suggest that hormones can induce apoptosis of bone tissue cells by activating the endoplasmic reticulum stress-PERK pathway, while ATG5-siRNA can reduce cell apoptosis by inhibiting the activity of this pathway (Fig. 7-b, e; Fig. 8 a-d).

### Core regulatory role and temporal dynamics of the endoplasmic reticulum stress-PERK pathway

Through temporal dynamic analysis of various indices, this study clarified the core regulatory role of the endoplasmic reticulum stress-PERK pathway in hormone-induced bone damage. The results showed that in Group B (hormone group), the expression of PERK, autophagy markers (e.g., ATG5, LC3), and apoptosis factors (Caspase family members) increased synchronously, showing consistency in temporal trends: all slightly increased at 1 week, sharply increased at 2 weeks, peaked at 3-4 weeks, when bone damage was most severe. In Group C (hormone + ATG5-siRNA group), intervention with ATG5-siRNA synchronously inhibited the expression of PERK and related molecules; all indices at each time point were significantly lower than those in Group B, and the degree of bone damage was also significantly alleviated. These temporal dynamic changes indicate that the PERK pathway plays a core regulatory role in the hormone-induced "endoplasmic reticulum stress-autophagy-apoptosis" cascade reaction, and inhibiting ATG5 expression via ATG5-siRNA can block this cascade reaction, providing a new strategy to antagonize hormone-induced bone toxicity (Fig. 7-e; Fig. 8 a-d).

## 4. Discussion

The widespread use of hormones has led to an annual increase in the incidence of steroid-induced osteonecrosis of the femoral head (SONFH) [20]. Studies indicate that SONFH pathogenesis involves cellular autophagy, with steroids regulating factors like Beclin-1 and LC3 [21]. Glucocorticoid (GC)-induced osteoblast apoptosis constitutes its biological basis [22], and endoplasmic reticulum stress (ERS)-mediated autophagy and apoptosis contribute to its development and treatment [23]. However, the interplay between autophagy and apoptosis, and the regulatory role of ERS in SONFH, remain incompletely understood.To investigate the associations among osteocyte autophagy, apoptosis, and SONFH, the regulatory mechanism of the ERS signaling pathway on these processes, and the inhibitory role of ATG5-siRNA, this study established three groups: control group (Group A), methylprednisolone group (Group B), and methylprednisolone + ATG5-siRNA group (Group C). The mRNA and protein expression of autophagy-related markers (Beclin1, MAP1LC3, ATG5) and the ERS sensor PERK in rat femoral heads were assessed to provide insights into SONFH pathogenesis, prevention, and treatment.

The main animal model findings were: (1) GCs promote osteocyte autophagy and ERS. (2) GCs enhance osteocyte apoptosis. (3) ATG5 induces osteocyte apoptosis via the ERS pathway. (4) ATG5 inhibitors alleviate osteocyte apoptosis by blocking the PERK branch of ERS. Under physiological conditions, autophagy promotes cell survival. However, under stimuli like hormones or hypoxia, excessive autophagy leads to organelle and cytoplasmic degradation, inducing cell death [24]. She et al. [25] found elevated expression of autophagy-related genes in osteoblasts from SONFH necrotic bone versus normal tissue. Blocking the AMPK/mTOR pathway reduced osteoblast autophagy. Li et al. [26] similarly concluded that regulating osteoblast autophagy represents a potential SONFH therapeutic target. Beclin-1 initiates autophagy and participates in autophagosome formation [27]. LC3 is critical for autophagosome formation and maturation. LC3-I conjugates with ATG7 and ATG3 to form LC3-II. The ATG5-ATG12 complex positively regulates this conjugation, underscoring ATG5’s crucial role in autophagy [28–30]. Therefore, Beclin-1, MAP1LC3, and ATG5 reflect cellular autophagy levels and autophagosome activity. Transmission electron microscopy revealed significantly more autophagosomes in the methylprednisolone group (Group B) than in the control group (Group A). PCR and Western blotting showed significantly higher mRNA and protein expression of Beclin-1, MAP1LC3, and ATG5 in Group B versus Group A, confirming methylprednisolone promotes osteocyte autophagy. HE staining revealed empty bone lacunae, fractured and disordered trabeculae in Group B. Scanning electron microscopy showed damaged osteocyte structure, further confirming methylprednisolone promotes osteocyte autophagy and death.

Endoplasmic reticulum stress (ERS) is a cellular response to stimuli that maintains protein homeostasis. Upon misfolded protein accumulation exceeding folding capacity, ERS triggers the unfolded protein response (UPR). If damage is irreparable, ERS induces apoptosis [31–32]. The ER regulates protein synthesis and folding by activating three pathways—IRE1α, PERK, and ATF6—to maintain homeostasis. Interactions among these pathways promote cell survival [33–35]. As PERK is a major ERS pathway and key mediator of apoptosis [36], it served as the detection target in this experiment. PCR and Western blotting showed higher PERK protein expression in Group B (methylprednisolone) than in Group A (control), indicating GCs promote ERS via PERK. Compared to Group B, mRNA and protein expression of autophagy markers (Beclin1, MAP1LC3, ATG5) and PERK were decreased in Group C (methylprednisolone + ATG5-siRNA), indicating ATG5-siRNA inhibits GC-induced osteocyte autophagy and ERS. Studies demonstrate ERS is associated with SONFH pathogenesis, involving osteocyte autophagy, apoptosis, and vascular injury [37]. Zheng et al. [38] found ATG5 regulates autophagy, ERS, and apoptosis via PERK, consistent with our findings.

Through integrated bioinformatics analysis, cellular experiments, and animal studies, this work systematically elucidates core pathological mechanisms and therapeutic targets in steroid-induced osteonecrosis of the femoral head (SONFH).GEO transcriptome analysis identified significant gene expression alterations in SONFH versus controls. Key differentially expressed genes (DEGs)—including BECN1, CASP3, ATG5, and EIF2AK3—showed enrichment in autophagy, apoptosis, and ER stress pathways. These genes further participate in dysregulated osteogenesis, collagen disorganization, and immune activation. Cellular assays confirmed hormonal activation of autophagy through ATG5-dependent pathways. Crucially, ATG5 forms extensive connections with ER stress mediator EIF2AK3 (PERK) and apoptotic executors (CASP3/9/12). These components establish a positive feedback loop driving apoptosis. In vivo validation demonstrated that glucocorticoid exposure triggers excessive autophagy progressing to autophagic death, while concurrently inducing apoptosis via ER stress-PERK signaling. This cascade resulted in trabecular fractures and elevated empty osteocyte lacunae. Collectively, dysregulation of the autophagy-apoptosis-ER stress axis—combined with impaired bone matrix metabolism and immune activation—underlies SONFH pathogenesis. As the central network node interconnecting these processes, ATG5 represents a promising therapeutic target for SONFH intervention(Figure 9).

**Fig 9.**
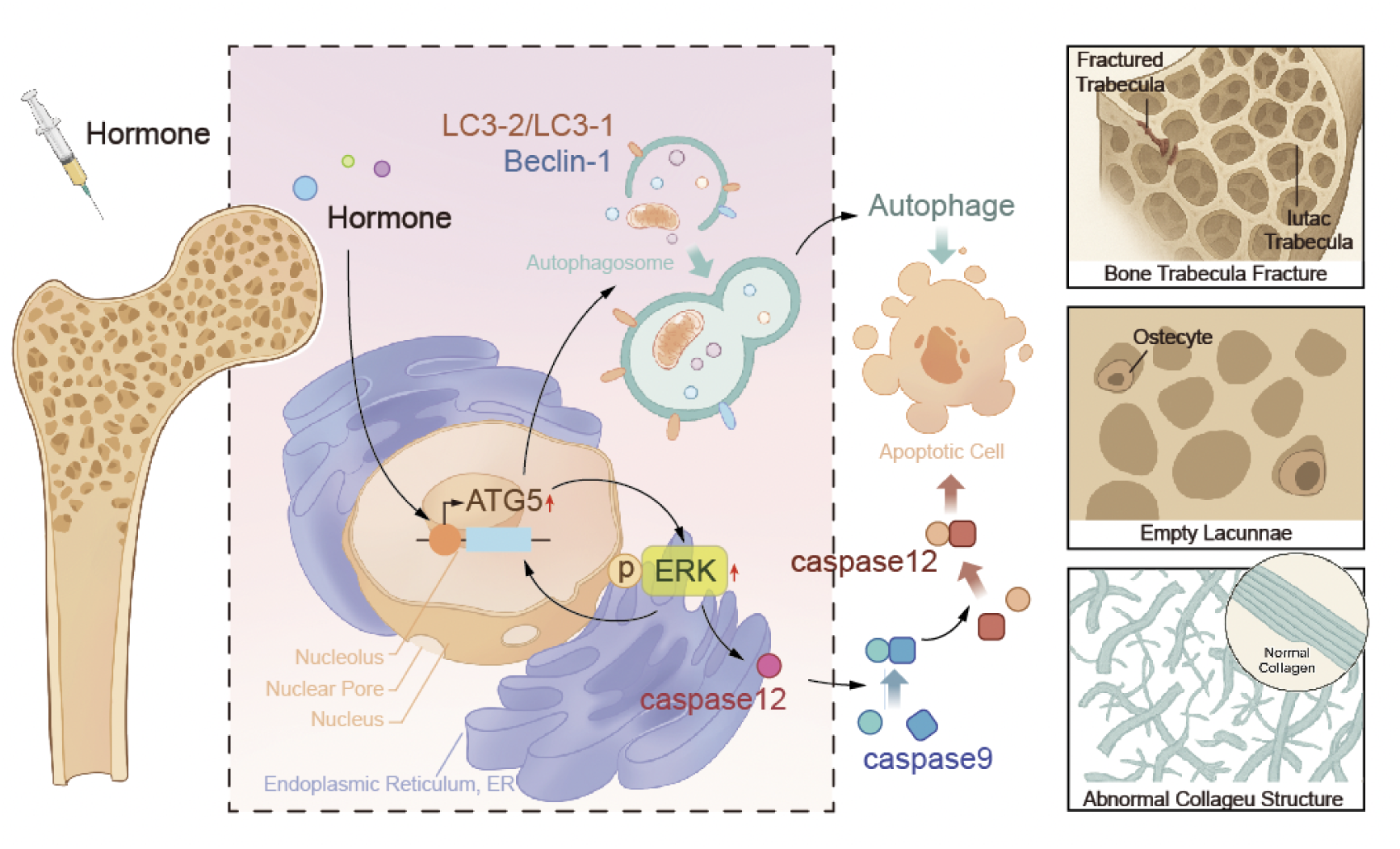
Schematic diagram of the mechanistic hypothesis

## Funding

The study was supported by grants from the National Natural Science Foundation of China (8196090146 ).

## Ethics approval and consent to participate

The animal study protocol was approved by the Medical Ethics Committee of Inner Mongolia Medical University(protocol code YKD202101138; approval2021/3/3).The study was conducted in accordance with internationally recognized principles for the use and care of laboratory animals.

## Data Availability Statement

The analyzed datasets generated during the study are available from the corresponding author on reasonable request.

## Acknowledgments

We would like to thank our colleagues for their useful discussions and comments and the support of funding project.

## Conflicts of Interest

The authors report no conflicts of interest with this study.

## Additional information

None.

## Author Contributions

Kunkun Liu: Methodology, Investigation, Writing-original draft, Writing-review & editing, Final approval. Boyong Jiang: Conceptualization, Investigation, Writing-review & editing, Final approval. Wanlin Liu: Investigation, Writing-review & editing, Final approval. Yulin Gong: Conceptualization, Writing-original draft, Writing-review & editing, Final approval. Zhenqun Zhao: Conceptualization, Writing-original draft, Writing- review & editing, Final approval, Funding acquisition.

